# Interpreting biochemical text with language models: a machine learning framework for reaction extraction and cheminformatic validation

**DOI:** 10.1101/2025.05.15.654376

**Authors:** Daven Lim, Swathi Badrinarayanan, Kira Sterling, Guru Rajesh, Eshaan Mistry, Daphne Yang, Max Lee, Kenneth Bryan Hsu, Mrunali Manjrekar, Cassandra Areff, Phil Xie, Ivan Kristanto, Arjun Chandran, J. Christopher Anderson

## Abstract

Recent advancements in large language models (LLMs) offer new opportunities for automating the manual curation of biochemical reaction databases from scientific literature. In this study, we present an integrated pipeline that enhances LLM-based extraction of enzymatic reactions with machine learning and cheminformatics-informed validation. Using BRENDA-linked PubMed articles, we evaluate GPT-4’s ability to extract reactions and infer missing chemical entities in textual descriptions of enzymatic reactions. Extracted reactions are converted to SMILES and InChI notations before being encoded into molecular fingerprint similarity scores and atom mapping metrics. These cheminformatics metrics are then used to train machine learning classifiers that validate GPT extractions. We employ a Positive-Unlabeled learning approach with synthetic invalid reactions to train various classifiers and assess model performances. The best classifier is then benchmarked on GPT extractions. Our findings show that GPT can accurately infer incomplete reactions and cheminformatics tools can serve as effective predictors of reaction validity. This work demonstrates a scalable framework for automated and reliable curation of enzymatic reaction databases, highlighting the potential of combining LLMs with cheminformatics and machine learning for reliable scientific knowledge extraction.

**Author Summary:** Curating databases of biochemical reactions is a time-consuming and manual task, yet it plays a vital role in advancing research in biology and chemistry. Many scientific articles describe important enzymatic reactions, but often do so in incomplete ways—such as mentioning only the starting molecule or the enzyme, and leaving out the rest. In this work, we explore how recent advancements in artificial intelligence, specifically large language models like GPT, can help extract such information automatically from scientific literature. We show that these models can not only find reactions in text, but also infer missing parts of reactions based on the surrounding context. To make sure these inferred reactions are chemically plausible, we use computational chemistry tools that analyze the structure of the molecules involved. We then train a machine learning model to help us automatically detect which reactions are likely to be valid. This combination of tools offers a new way to speed up and improve how biochemical knowledge is extracted from the growing body of scientific literature. Our study suggests that this kind of automation could help scientists keep biological databases up to date and reduce the burden of manual data entry.

## 1 Introduction

The constant update of chemical reaction databases from scientific literature is essential for the advancement of computational chemistry and biochemistry. Despite a large volume of open-access reaction databases available — such as BRENDA [1], Kegg [2], Metacyc [3], Open Reaction Database (ORD) [4] and Chemical Reaction Database (CRD) [5] — the extraction of reactions from academic literature has largely relied on manual curation. This process is time-consuming and can result in databases that are incomplete, unstandardized, and occasionally contain erroneous entries.

Advancements in natural language processing (NLP) techniques and large language models (LLMs) have enabled the design of effective tools to automate extraction of scientific information from text. For example, fine-tuning pre-trained LLMs such as GPT-3 and Llama-2 has been demonstrated to be effective for the extraction of records from scientific text in structured formats [6]. In addition, guiding GPT-3.5 and GPT-4 with prompt engineering and fine-tuning GPT-3.5 and other LLMs such as Llama2 and T5 have also been shown to achieve state-of-the-art performance on the task of chemical text mining [7].

In the context of chemical reaction mining, LLMs such as GPT-3.5 and Gemini 1.0 Pro have been shown to be able to extract chemical reactants and products efficiently from patent documents from the USPTO (US Patent and Trademark Office) [8]. However, a key challenge in applying transformer-based LLMs to expertise-intensive domains like chemistry is its potential to make mistakes [8]. In particular, the topic of processing reactions post-extraction is still one for which limited research has been published.

Extracted reactions can occasionally be incorrect, a limitation we sought to address by investigating automated methods to validate and correct these outputs. In addition, we observe that scientific literature on enzymes often includes incomplete reactions, most often only explicitly mentioning substrates and the type of reaction it undergoes by name while leaving products unreferenced. This presents a challenge to automation of reaction extraction but is particularly common in enzymatic reaction papers, many of which are curated into the BRENDA database, thereby posing a significant challenge to full automation.

Therefore, our goal of automating curation of reaction databases via extraction from scientific literature goes beyond merely identifying reactions that are mentioned in text. An understanding of the chemical context discussed in a paper is needed for accurate and comprehensive extraction—particularly when reactions are often incompletely referenced in the literature. LLMs must still be capable of identifying reaction components even when parts of reactions such as products may be unspecified. In such cases, the ability of LLMs to infer missing components based on context, such as reaction type or enzyme function, becomes significant. This highlights the importance of an understanding beyond textual parsing in enabling comprehensive extraction of reaction data.

In this work, we present methods to supplement LLM-based automated enzymatic reaction extraction. To address incomplete in-text references of reactions, we evaluate the ability of LLMs to not only extract enzymatic reactions but also infer missing chemical entities to enhance the completeness of reaction curation. On post-extraction processing, we utilize cheminformatics tools such as molecular fingerprints and atom mapping to encode reactions and subsequently computationally validate the plausibility of LLM-extracted reactions via cheminformatics-based machine learning models. Fig 1 summarizes the pipeline in this study. Through these evaluations, we demonstrate the potential of these methods in augmenting automated LLM-based chemical reaction extraction.

**Fig 1:**
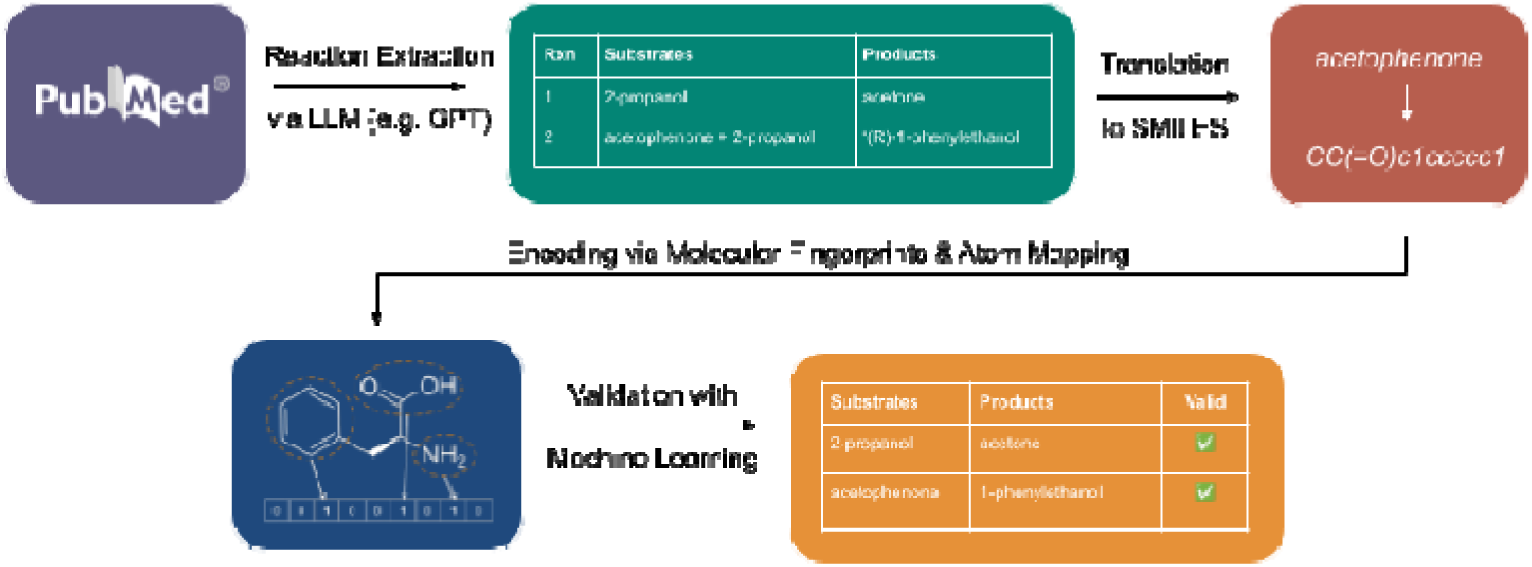
Pipeline for automated extraction and validation of chemical reactions. Extraction of biochemical reactions from scientific literature, such as papers found in PubMed, is performed via LLMs (e.g., GPT), generating substrate-product pairs and the enzymes associated with each reaction. These extracted reactions are then converted to SMILES (Simplified Molecular Input Line Entry System) notation. Molecular fingerprints similarity scores and atom mapping metrics are generated to encode the reactions into features. Machine learning models are trained on these features in order to validate extracted reactions. Validated reactions can then be used to populate a comprehensive reaction database.

## 2 Results

### 2.1 GPT Extraction

To demonstrate the challenge of automated reaction extraction with incomplete reaction references in text and subsequently evaluate the performance of LLMs in inferring unreferenced reaction products, we use GPT-4o-standard to perform the task in this study. We evaluate the model on its performance in extracting enzymatic reactions and inferring missing chemical entities in incompletely referenced reactions. A prompt was passed into GPT as input, along with the full text of a scientific research paper.

Comparing GPT’s extractions with entries in the BRENDA database, GPT has demonstrated to perform well in the extraction-only task, with it being able to identify many reactions that BRENDA has curated. All reactions that GPT had extracted can be found in the paper manually. Of the reactions listed in BRENDA but not extracted by GPT, many of them are found in figures, suggesting that GPT struggles to interpret compounds referenced in figures. This is also observed in prior studies with LLMs [8]. Of other reactions that GPT misses, some could not be found through manually looking for them in the corresponding papers to which they are linked. In addition, GPT was also observed to be able to identify reactions that are present in the literature but are missed by BRENDA, suggesting that GPT can effectively complement BRENDA. Specific extractions on a subset of 22 papers and their respective BRENDA entries are included in supplementary material.

Additionally, papers linked to BRENDA often reference reactions only partially. A common pattern is that substrates are mentioned without corresponding products, such as in the sentence: “… efficiently catalyze the oxidation of primary alcohols such as 1-hexanol and 1-butanol but not secondary alcohols such as 2-propanol”. In a subset of 100 such papers, more than 75% of the reactions that were missed by GPT but listed in BRENDA were only partially referenced in literature, such as having products left out. GPT was then prompted to extract these reactions and complete them. The validity of these extracted reactions was then evaluated using cheminformatics-based machine learning validation.

### 2.2 Molecular Fingerprints

Molecular similarity between substrates and products can be used to assess the validity of reactions extracted by GPT, as a low average similarity score across a reaction may indicate that a reaction is implausible and thus incorrectly extracted, due to large changes in its chemical composition. To evaluate this, we used RDKit (version 2022.09.5) to generate molecular fingerprints, encoding each molecular structure into a machine-readable numerical format, typically represented as a bit or integer vector or string. These fingerprints allow for efficient representation of a molecule’s structure and enable convenient comparisons. In this study, we incorporated the three major classes of molecular fingerprints [9] — substructure key-based, topological or path-based, and circular fingerprints. Substructure key-based fingerprints, such as MACCS (Molecular ACCess System) Key Fingerprints, encode the presence of predefined substructures such as functional groups within a molecule. Topological or path-based fingerprints, including Hashed Topological Fingerprints and Atom Pair Fingerprints, capture information about atomic connectivity along molecular paths. Circular fingerprints, notably the Extended Connectivity Fingerprint (ECFP), also known as a Morgan fingerprint in RDKit, represent the atomic environment around each atom within a specified radius.

In our model, these fingerprints were implemented and compared to evaluate the plausibility of enzymatic reactions. Specifically, MACCS keys fingerprints were used to detect predefined substructures, Hashed Atom Pair fingerprints encoded molecular structures by hashing paths between atom pairs, and ECFP (Morgan fingerprints) captured atomic neighborhoods within a set radius. The similarity between the reactants and products of each reaction was then calculated using the Tanimoto coefficient, a standard similarity metric in cheminformatics [10].

### 2.3 Atom Mapping

Atom mapping identifies correspondences between atoms in reactants and those in products, enabling detailed mechanistic analysis of chemical reactions. This mapping is critical for downstream tasks such as mechanistic inference. Indigo and RXNMapper are two popular atom mapping tools, which are appropriate for this task based on a benchmarking comparison of atom-mapping tools [11]. These tools represent two distinct methodological approaches. Indigo uses subgraph matching to find a one-to-one correspondence between atoms by identifying reaction centers. RXNMapper [12], on the other hand, is a transformer neural network model. RXNMapper was trained to align atoms by using unsupervised attention mechanisms without requiring hand-labeled mappings, and has been shown to outperform traditional rule-based approaches, especially on complex or noisy reaction datasets [11]. In this study, we utilize RXNMapper to perform atom mapping.

To evaluate mapping quality, atom mapping consistency metrics were computed with a focus on the preservation of the carbon backbone, a fundamental structural feature in organic molecules. Assessing how well this backbone is conserved serves as a proxy for reaction plausibility, as enzymatic transformations often preserve the carbon framework, while often involving substitutions, additions, or reductions that occur at functional groups rather than modifying the core carbon skeleton. Therefore, atom mapping metrics that quantify the degree of backbone rearrangement or fragmentation provide insight into the plausibility of the extracted reaction. In addition, unmapped carbon atoms often indicate failures associated with mapping, which serves as a key feature in identifying invalid chemical transformations. Fig 2 shows an example of a mapped reaction by RXNMapper and the corresponding metrics generated.

**Fig 2.**
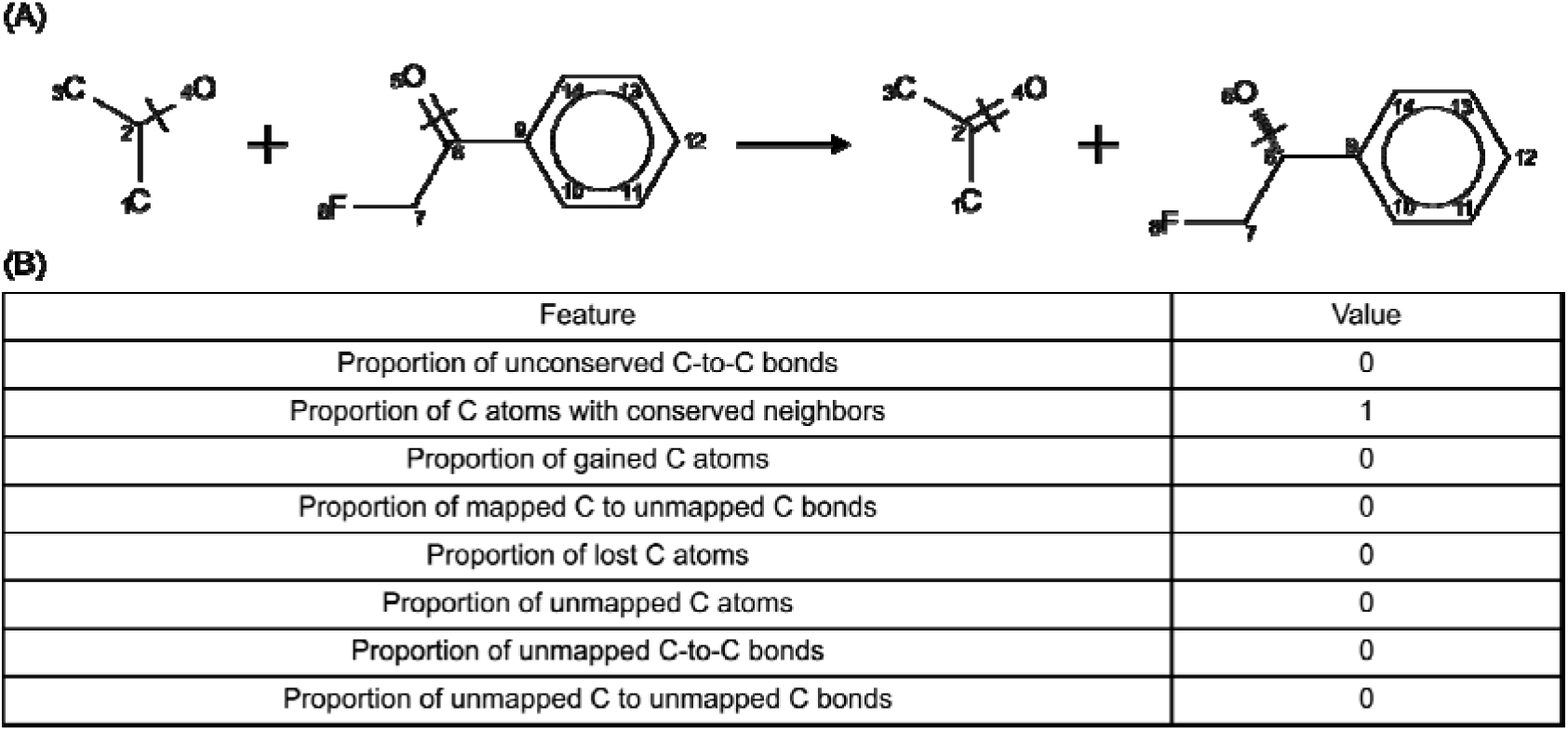
Atom mapping and mapping-based metrics for a chemical reaction. (A) Atom-level mapping of a chemical reaction generated using RXNMapper, with atoms numbered to indicate correspondence between reactants and products. (B) Quantitative metrics summarizing structural conservation and atom mapping consistency for the reaction shown.

### 2.4 Machine Learning Approach for Reaction Validation

To validate reactions extracted by GPT, we trained machine learning models using the cheminformatics features — namely, molecular fingerprint similarity scores and atom mapping metrics. These models classify reactions as valid or invalid based on those features. We used a random forest classifier due to its interpretability and robustness, in addition to other models such as logistic regression, naïve Bayes, and support vector machines. Each model was trained on 430 valid and 430 synthetic invalid reactions with an 80-20 train-validation split using 5-fold cross validation. Key performance metrics, including accuracy, precision, recall, and F1-score, were reported. We also assessed feature importance using the mean decrease in impurity in the random forest.

The approach described aligns with a form of Positive-Unlabeled (PU) learning, which is a type of semi-supervised learning where only positive examples and unlabeled data (which may contain both positive and negative examples) are available [13]. While curated databases like BRENDA provide positive examples, there are no large-scale, manually labeled invalid reaction datasets. PU learning assumes that negative examples differ in distribution from positive ones. In the first step, reliable negatives are identified from the unlabeled set. In the second, these are used along with positive examples to train a supervised classifier. We followed this approach and evaluated model performance via 5-fold cross-validation, comparing the random forest against the other model types.

### 2.5 Synthetic Invalid Reactions

Positive examples were sourced from BRENDA. To construct the synthetic negative set for PU learning, we permuted substrate-product pairs from valid BRENDA reactions, ensuring that the substrates and products each originated from different original reactions, and manually removed reactions that closely resemble valid reactions. Fig 3 shows the distribution of a subset of features across the two datasets. We can observe differing feature distributions across both sets of reactions.

**Fig 3.**
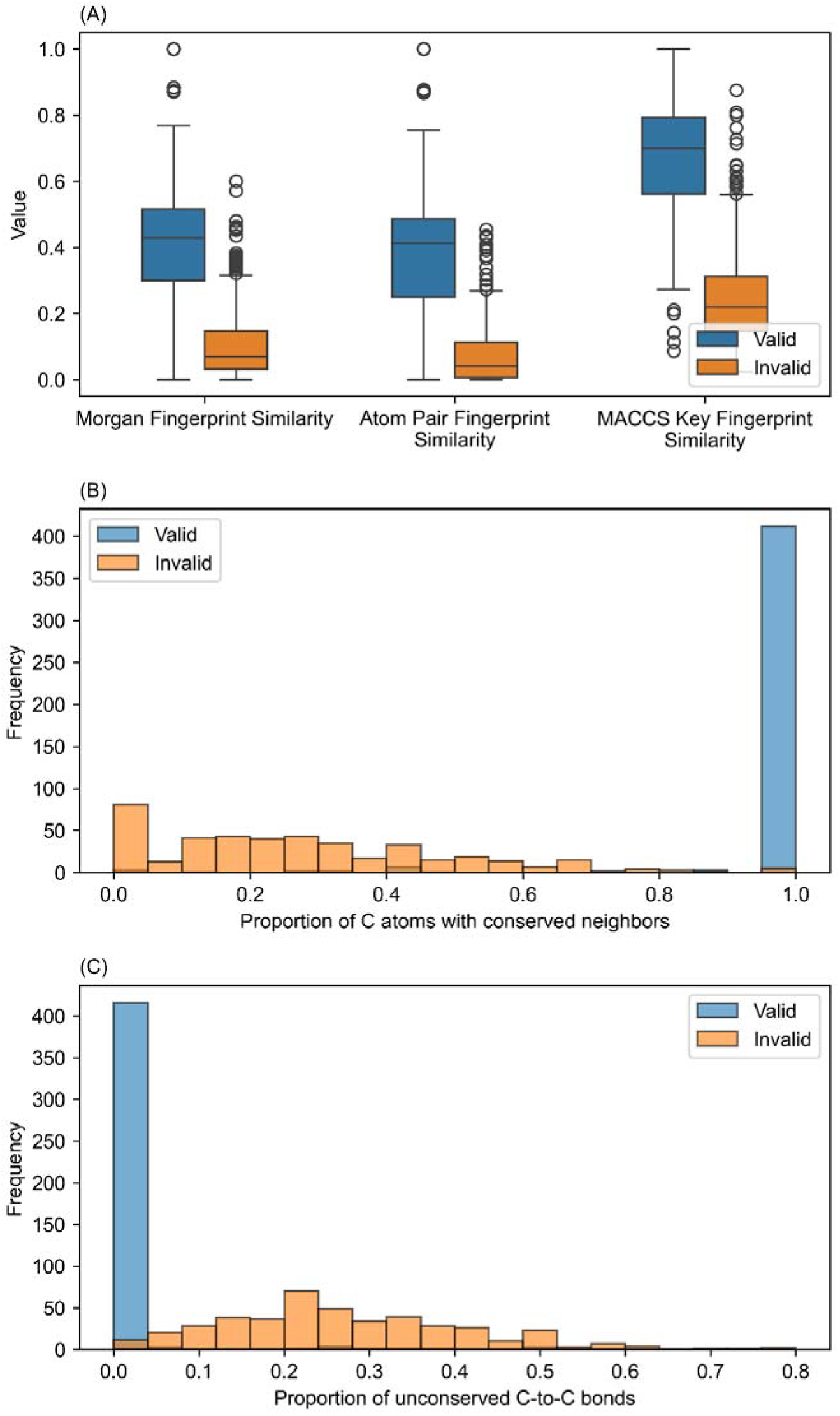
Feature distribution comparison between valid and synthetic invalid reactions. (A) Average molecular fingerprint similarity across multiple fingerprint types. (B) Proportion of carbon atoms with conserved neighboring atoms. (C) Proportion of unconserved carbon–carbon bonds.

### 2.6 Machine Learning Models

The model functions as a first layer of automated screening for GPT’s extractions. It uses cheminformatics metrics, in particular molecular fingerprint similarity across reactions and atom mapping metrics, to assess validity of the extracted reactions to account for potential errors or hallucinations from GPT. Reactions classified as potentially invalid will be flagged for review, allowing for expert verification on a significantly smaller subset of extracted reactions. This can significantly enhance accurate and reliable automated curation of reaction databases using LLMs.

We first benchmarked our model’s performance using 5-fold cross validation on our dataset, where Fig 4A shows the performance of the various machine learning models. Models trained include random forest, logistic regression, naive Bayes, and support vector machine, where the random forest outperforms the other models in almost every metric. Recall measures the model’s ability to correctly identify negative examples from all actual negatives in the dataset, and the random forest model is observed to exhibit consistently high recall. This is significant, because while misclassifying valid reactions as invalid can be corrected during post-validation review, the reverse—allowing invalid reactions to pass as valid—poses a greater risk, as it directly impacts the reliability of the curated data.

**Fig 4.**
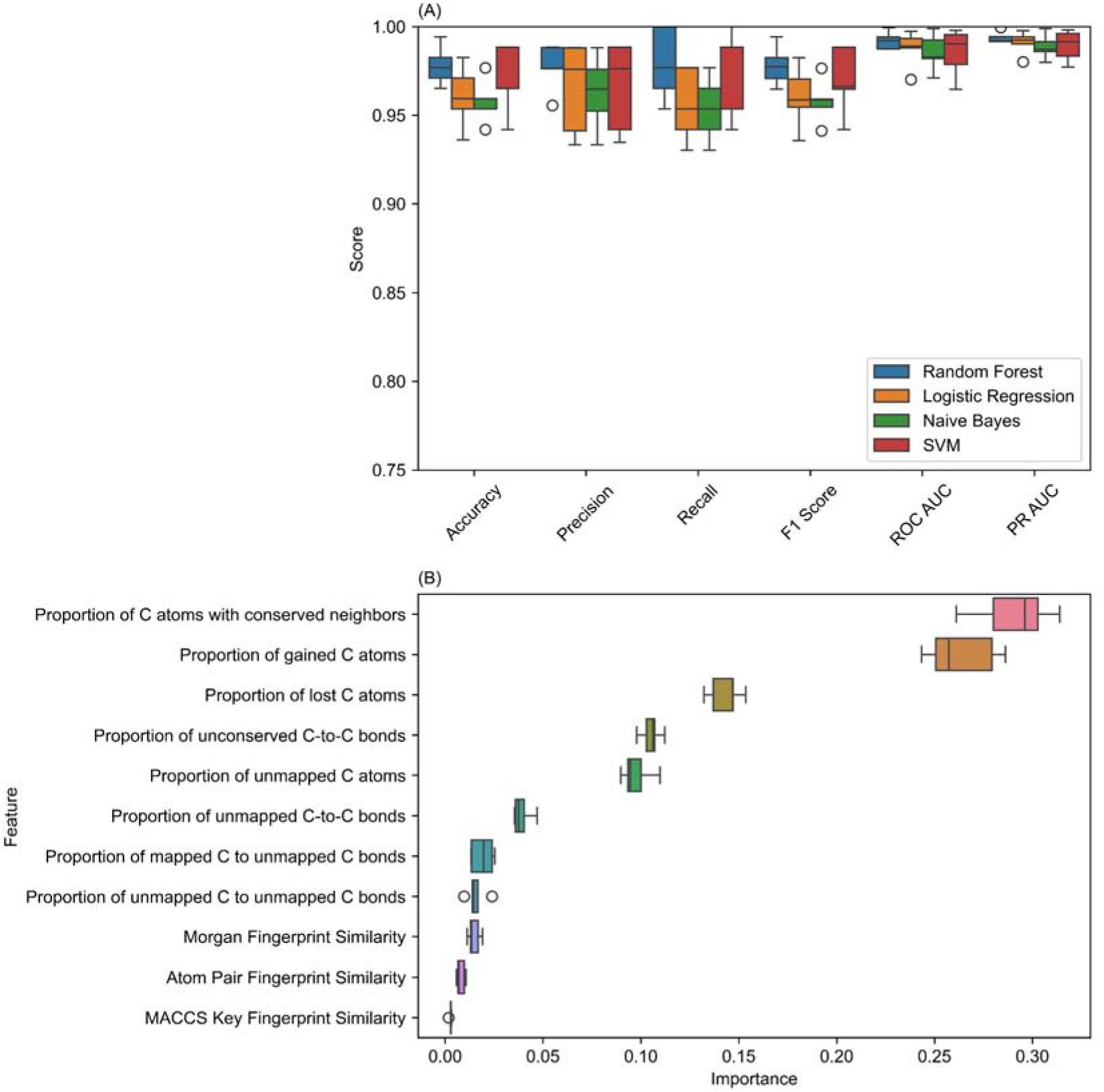
Machine learning model performance and feature importance for reaction validation. (A) Cross-validation results comparing the performance of four classifiers—random forest, logistic regression, naive Bayes, and support vector machine—trained on a synthetic invalid reaction dataset. The random forest classifier achieves the highest recall, favoring reliable identification of invalid reactions. (B) Feature importance scores for atom mapping and molecular fingerprint similarity metrics, indicating the significance of carbon backbone conservation in predicting reaction validity.

Atom mapping metrics used in our model reflect the degree to which the carbon backbone is preserved from reactants to products. In our results, atom mapping features, particularly those measuring conserved carbon–carbon bonds and neighborhood preservation, were found to contribute most heavily to the model’s predictive performance (see Fig 4B). This suggests that the preservation of the carbon backbone is a strong predictor of reaction validity.

That said, both atom mapping and molecular fingerprint metrics carry their own predictive utility. Fingerprint similarity captures broader chemical resemblance between reactants and products, which can be useful for detecting huge chemical transformations. When combined, these two types of cheminformatics features provide complementary perspectives, with one focused on structural correspondence and the other on molecular similarity, which offers a robust basis for validating LLM-extracted reactions.

### 2.7 Performance of GPT Extraction on Incomplete Reactions

While tools like Molecular Transformer [14] have shown strong performance in predicting reaction products from known reactants and reagents, our study investigates whether general-purpose LLMs trained primarily on textual data can also infer products based on textual descriptions of the substrate and reaction type (e.g., oxidation). We find that GPT is capable of performing this task to a large extent.

To evaluate its performance, the extracted reactions are first compared against entries in the BRENDA database, which are assumed to be valid. We compared the reactions using InChI [15] notation for automation and standardization. A substantial number of GPT-extracted reactions match those in BRENDA. The remaining reactions, which are not found in BRENDA, are manually reviewed by consulting literature. This manual verification confirms that all GPT-extracted reactions are valid.

Next, GPT-extracted reactions are first into cheminformatics features before testing the trained model on these reactions. Fig 5 shows the distribution of the model’s validations on GPT-extracted reactions. A large proportion of extracted reactions are classified as valid, indicating that the model can effectively identify valid reactions. There is a small proportion of reactions that are classified as potentially invalid. Looking into this further, these reactions often involve multi-step transformations or large structural changes. An example is acetyl CoA reduction to ethanol in a two-step reaction, shown in Fig 6A. Such transformations are valid but are also less likely to preserve the carbon backbone or functional groups and are harder to assess without mechanistic insight. In these cases, more specialized tools with mechanistic insight may be needed to complement the current features to improve classification accuracy. Other reactions that are classified as invalid include reactions where more substrates than products are extracted, such as the reaction shown in Fig 6B, which could lead to atom mapping metrics reflecting a larger number of unmapped carbon atoms. Examples of reactions and their classifications by the model and by manual verification are provided in the supplementary material, drawn from the same subset of 22 papers referenced earlier for GPT’s extractions. As discussed earlier, misclassifying valid reactions as invalid can be corrected with minimal impact.

**Fig 5.**
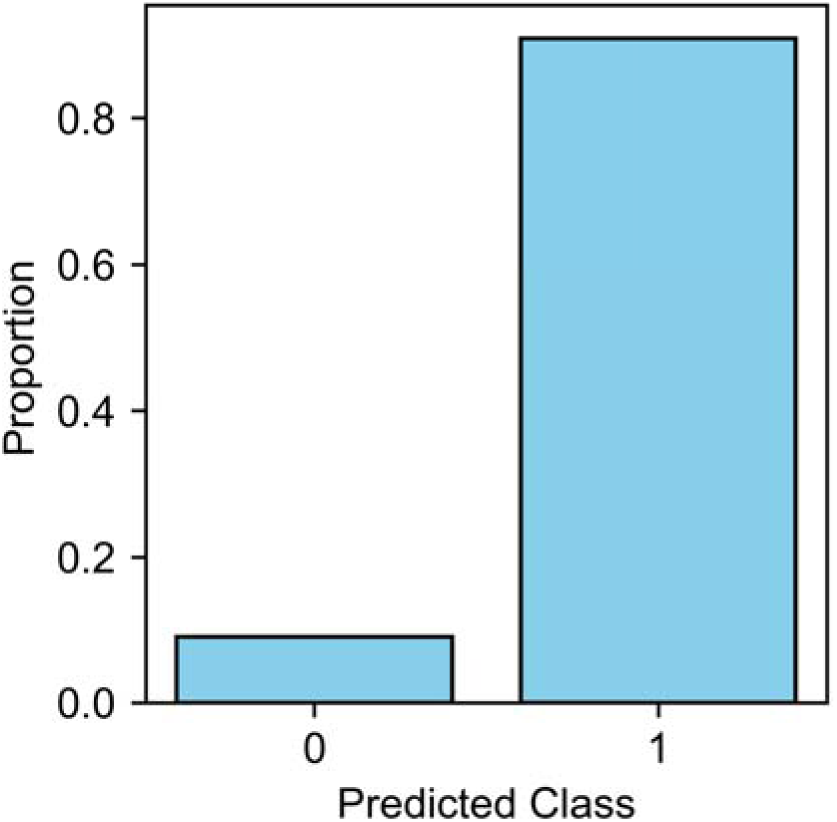
Validation outcomes of GPT-extracted enzymatic reactions using a machine learning classifier.

**Fig 6.**
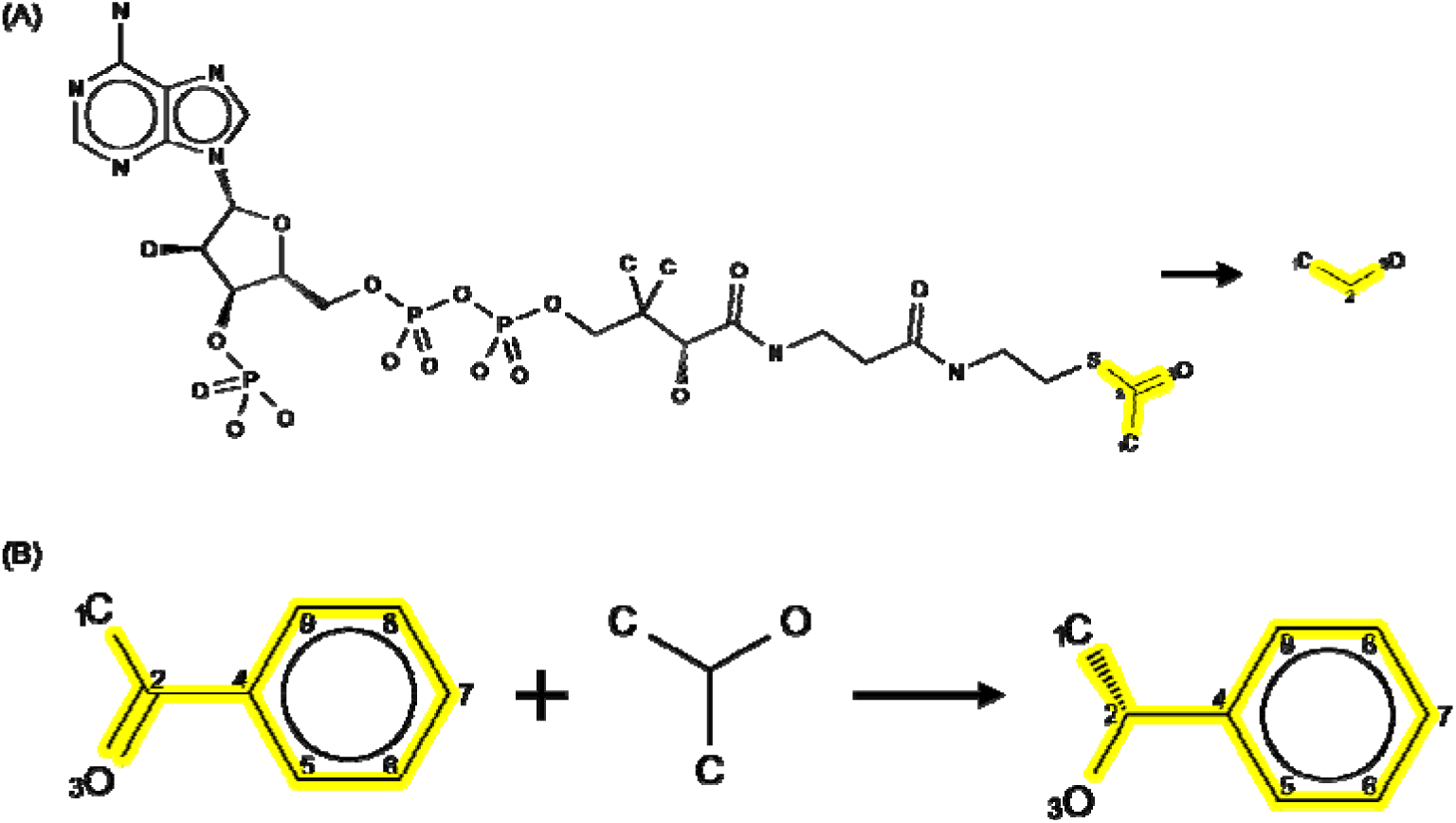
Examples of GPT-extracted reactions classified as invalid by the model. (A) A two-step transformation from acetyl-CoA to ethanol is flagged due to low carbon backbone conservation, despite being chemically valid. (B) An imbalanced reaction with more substrates than products results in poor atom mapping metrics, leading to a false classification as invalid.

We also observed that comparing InChI identifiers, rather than SMILES [16], significantly improved matching with BRENDA. Still, stereochemical variants (e.g., cis/trans or E/Z isomers) could yield different InChI strings for the same named molecule, requiring downstream validation via cheminformatics models. Discrepancies between GPT extractions and BRENDA entries were often due to differences in chemical naming, such as abbreviations (e.g., “KDG” vs. “2-keto-3-deoxy-D-gluconate”), or inconsistent inclusion of stereochemistry, though the underlying reaction being described remained the same. Here, while comparing InChI names of extracted reactions to those in BRENDA would not have validated these extracted reactions, they will be validated by our model to verify that these are plausible reactions.

## 3 Discussion

This study introduces a framework for enhancing automated reaction extraction from scientific literature by integrating large language models (LLMs) with cheminformatics and machine learning techniques. We demonstrate that cheminformatics tools, such as molecular fingerprints and atom mapping, are effective for validating the plausibility of extracted reactions. Features like the preservation of carbon backbones serve as strong predictors of enzymatic reaction validity, providing a basis for automated plausibility screening.

In addition, GPT has shown strong capability in inferring missing components of incompletely referenced reactions, highlighting the potential of even general-purpose LLMs for chemical information extraction even without domain-specific fine-tuning. This expands the practical utility of LLMs in biochemistry and reaction curation pipelines. Future work may focus on fine-tuning LLMs specifically for chemical and biochemical text to enhance recognition of chemical entities and reaction context. It is also important to note that GPT’s predictions may be biased toward reactions frequently encountered during training, and future work can explore its performance in generalizing to novel or less-represented reactions beyond its training distribution.

Additionally, while cheminformatics tools used in this study performed well, they may fall short in capturing more complex or nuanced chemical transformations, such as in multi-step reactions. Incorporating tools that offer mechanistic insights can improve validation performance, and benchmarking the cheminformatics tools used in this study with more mechanistically-involved tools can quantify potential performance improvements. Further work can also look into the generation of synthetic invalid reactions that can help machine learning models learn more subtle features of plausible reactions, beyond focusing on carbon backbones and main functional groups. With the demonstrated effectiveness of combining LLMs, cheminformatics, and machine learning, the path toward scalable, automated, and accurate reaction database curation appears highly promising.

## 4 Materials and Methods

### 4.1 LLMs

A prompt was passed into GPT using OpenAI’s API as input along with the full text of a scientific research paper, with different prompt evolutions evaluated at their ability to maximize extraction recall. Prompts were engineered and improved using the Automatic Prompt Engineer (APE) method [17], and few shot prompting was utilized to provide examples of extractions of enzymatic reactions from text. We tailored our prompts to address incomplete reactions by prompting GPT to complete these partially referenced reactions based on its knowledge of chemistry. The final prompt we used is included in the supplementary material.

### 4.2 Dataset

BRENDA (BRaunschweig ENzyme DAtabase) is a comprehensive relational database on enzymes and their reactions, curated based on primary literature linked to PubMed from over 150,000 literature references [1]. The enzyme-catalyzed reactions in BRENDA are stored alongside PubMed references to the literature they are sourced from, which enabled us to apply large-language models (LLMs) like GPT-4 to the same papers examined by BRENDA contributors. We leveraged this correspondence to compare the performance of LLMs with BRENDA on the task of chemical reaction extraction on the same corpus of scientific literature, and identify areas for potential augmentations in LLM-based extractions. To acquire the full articles to evaluate LLM performance on this task, we downloaded individual articles from the PubMed Central Open Access Subset. References in BRENDA are done using the PubMedID, which we linked to the PMC ID via the E-Utilities Service (Entrez Molecular Sequence Database System).

### 4.3 Conversion of Chemical Names to SMILES and InChI Formats

After extracting substrates and products from literature, chemical names are converted into SMILES notation to be passed as input to our following cheminformatic methods. To achieve this, three chemical databases were queried: NIH PubChem [18], NCI CACTUS Chemical Identifier Resolver [19], and ChemSpider [20]. Of the 1700 chemical names queried, these databases returned an equivalent SMILES entry about 90% of the time. Though these three databases contain a multitude of equivalent synonyms for each unique molecule, the remaining 10% of chemicals not found can be attributed to different or non-standard naming conventions, or that the chemical was simply not available in the databases. Human-in-the-loop resolutions may be needed for these edge cases.

The SMILES notations are then converted into InChI (IUPAC International Chemical Identifier) using RDKit to compare GPT extractions with the entries in BRENDA. While one molecule could have multiple SMILES representations, InChI representations are unique to a chemical structure and provide a standardized format that enables comparison and captures equivalence between different chemical names. This facilitates automatic standardized comparisons.

## Supporting information

Reaction entries obtained from the BRENDA database

Input prompt to GPT for reaction extraction

GPT extracted reactions and BRENDA reaction entries on a subset of 22 papers

GPT extracted reactions with cheminformatics metrics and model validation scores

## Acknowledgements

This work was conducted by iGEM at Berkeley, an undergraduate research club at the University of California, Berkeley. We gratefully acknowledge the support of our club’s sponsors: Tetsuwan Scientific, Associated Students of the University of California, UC Berkeley Engineering Student Council, Botchan Lab, Rulison Family, Tonkomo LLC, Pande Family, Spiegel Family, Jacob Pruess, Kanjana Taedullayasatit, Stuart Russell, Sophie Hahn, Lin Tan, Geneious, Michaela Healton, and Shen & Namiko Chen. We would also like to acknowledge Aidan Wen, Vivian Lu, Janani Sriram, Ayati Sharma, Tara Pande, Sravya Basvapatri, Jesus Del Rio, Jierui Xu, Jacob Luo, Nilasha Krishnamurthy, Kathan Gandhi, Sina Ghandiana, Vanessa Anderson, and Shreya Mohantya, all of whom contributed to this work as undergraduate members of iGEM at Berkeley.

## Supplementary Materials

All code and data used for running experiments are available on a GitHub Repository with a DOI at: https://doi.org/10.5281/zenodo.15428043

**S1 BRENDA Entries.csv.** Reaction entries obtained from the BRENDA database

**S2 GPT Prompt.docx.** Input prompt to GPT for reaction extraction

**S3 GPT vs. Brenda Annotations.xlsx.** GPT extracted reactions and BRENDA reaction entries on a subset of 22 papers

**S4 GPT Extracted Reaction Validations.csv**. GPT extracted reactions with cheminformatics metrics and model validation scores

